# Host Specificity and Resistance of Cowpea and Common Bean to *Elsinoë* spp. isolated from cowpea

**DOI:** 10.64898/2026.10.11.755143

**Authors:** S.O. Oriama, Y. Masheti, C. Ndinya, J.F. Kelele, S. Chepkurui, S. Misango

## Abstract

Cowpea (*Vigna unguiculata* L. Walp.) is an important leguminous crop in Kenya, but its production is constrained by diseases that reduce yield and market quality. Scab disease, caused by fungi in the genus *Elsinoë*, is particularly damaging due to its effects on foliage, flowers, and pods. This study investigated cowpea scab under controlled conditions, focusing on pathogen identification, host specificity, and varietal response. The pathogen was isolated from infected cowpea plants, and its identity as an *Elsinoë* sp. distinct from *Elsinoë phaseoli* was confirmed based on previously established multi-locus DNA barcoding (ITS and TEF1-α) and phylogenetic data. Pathogenicity and cross-infectivity tests were conducted on three cowpea and three common bean (*Phaseolus vulgaris* L.) varieties. Kunde Tamu and Kunde Soko were highly susceptible, showing increasing disease severity across growth stages. By flowering, lesions had spread extensively on leaves and stems, and during pod filling, more than 90% of plant tissue was affected. Pods developed severe scab lesions, often becoming mummified and failing to fill grain (severity score 5). Kat Kunde exhibited partial tolerance, with low disease severity (≤2) during vegetative and flowering stages and minimal symptom progression. However, severity increased during pod filling, with pod lesions and occasional mummification, resulting in a final severity comparable to susceptible varieties, indicating stage-specific tolerance. All common bean varieties (CAL 194, CAL 143, and GLP 2) showed no scab symptoms, with severity scores remaining at 1, confirming absence of cross-infectivity. Typical scab symptoms in cowpea included cork-like lesions on leaves, stems, and pods, leading to flower abortion and pod mummification in susceptible varieties. Cross-infectivity tests confirmed that the isolate possesses strict host specificity to cowpea, remaining entirely distinct in host range from *Elsinoë phaseoli* which infects common bean. These findings confirm *Elsinoë* sp. as the causal agent of cowpea scab in Kenya, demonstrate strict host specificity, and identify tolerant germplasm for resistance breeding and improved disease management.

## INTRODUCTION

Cowpea (*Vigna unguiculata* (L.) Walp.) is an important leguminous crop widely cultivated in sub-Saharan Africa, including Kenya, where it plays a critical role in food and nutritional security. It is primarily consumed as a leafy vegetable and as grain, providing an affordable source of plant-based protein, vitamins, and micronutrients. In many rural and peri-urban households, cowpea leaves are classified among African Leafy Vegetables (ALVs) and contribute significantly to dietary diversity (Owade et al., 2020). The grain is also an essential protein source, often referred to as “poor man’s meat” due to its high nutritional value (Smartt, 1986). Despite its importance, cowpea production remains relatively low compared to demand, with global demand estimated at 5–6 million metric tonnes annually against a production of approximately 2 million metric tonnes (Kenya National Bureau of Statistics, 2025). In Kenya, production is largely dominated by smallholder farmers practicing subsistence agriculture, often intercropping cowpea with other ALVs and grain legumes (Owade et al., 2019).

Cowpea cultivation is constrained by several biotic stresses, particularly diseases that affect both yield and marketability. These include viral, bacterial, and fungal diseases such as Bean Common Mosaic Virus (BCMV), anthracnose caused by *Colletotrichum lindemuthianum*, Common Bacterial Blight (CBB) caused by *Xanthomonas* spp., and scab disease caused by species of the genus *Elsinoë*. Among these, cowpea scab is of increasing concern due to its impact on both the vegetative and reproductive parts of the plant. The disease manifests as small, whitish-grey corky lesions on leaves and pods, often surrounded by dark purple margins, accompanied by leaf puckering and distortion. Severe infections lead to deformation of leaves and pods, rendering the vegetable unmarketable and reducing consumer acceptance. In grain production systems, infection during the reproductive stage may result in flower abortion, reduced pod set, and ultimately significant yield losses.

Cowpea scab has previously been reported and studied (Emechebe, 1980; Mungo et al., 1998); however, historical limitations in pathogen identification and documentation have led to confusion and possible misidentification of the causal agents. The disease is caused by pathogenic fungi belonging to the order Myriangiales, genus *Elsinoë*, which are known to infect a wide range of host plants. Members of this genus are responsible for scab diseases characterized by sunken lesions and corky outgrowths on infected tissues (Fan et al., 2017). Although similar symptoms are observed across different legume hosts, emerging evidence suggests that *Elsinoë* species exhibit a high degree of host specificity, with limited cross-infectivity between pathogens infecting different plant genera (Fan et al., 2017; Jayawardena et al., 2014).

Field observations in Kenya, including demonstration trials at KALRO Kakamega, have highlighted the severity of cowpea scab under conducive conditions. In these trials, the disease caused extensive damage, with characteristic white lesions forming along leaf veins, leading to lamina folding. Pods were severely deformed, exhibiting curling and mummification due to coalescing lesions on pods and stems. These observations closely reflect farmer-reported experiences, emphasizing the economic significance of the disease in local production systems.

Despite its importance, there remains a limited understanding of the host specificity and resistance mechanisms of cowpea to *Elsinoë* spp., particularly in relation to closely related, co-cultivated legumes such as common bean (*Phaseolus vulgaris* L.). Common bean is a primary candidate for comparative study because its congeneric sister species, *Elsinoë phaseoli*, causes common bean scab (Masheti et al., 2024a). Because smallholders in Western Kenya frequently intercrop cowpea and common bean, investigating whether the cowpea-isolated *Elsinoë* pathogen can breach host barriers to infect *P. vulgaris* is highly critical for regional biosafety and disease containment.

This baseline study aims to investigate the host specificity of an *Elsinoë* sp. isolated from cowpea, and to evaluate the spill-over infection to common bean and the resistance of economically dominant cowpea and common bean varieties to this local isolate. This provides foundational insights to support sustainable crop diversification, intercropping management, and targeted breeding strategies.

## METHODOLOGY

Two sets of experiments were set up at KALRO Kakamega to isolate and test the infectivity of the isolated pathogen on both cowpea (*Vigna unguiculata*. L) and common beans (*Phaseolus vulgaris*. L). Three different varieties of cowpea that are commonly grown in the region were used in the experiment while three different varieties of common beans were used for this study.

For the choice of the common bean CAL 143 was included because of its resistance to *Elsinoë phaseoli* which is a congeneric species of the pathogen being investigated (Masheti et al., 2024a; Oriama et al., 2024). The other two varieties GLP2 and CAL194 are highly susceptible to the *Elsinoë phaseoli* pathogen.

### Experimental set up

#### Set one

Six planting pots were filled with sterile soil mixed with manure and sand at the ratio of 3:2:1 and planted with each variety so that each variety of cowpea and common bean had six pots and a total of 36 plants. DAP fertilizer was applied by placing a pinch of granules on the pot’s soil a week after germination. The plants were watered daily while monitoring the weeds growth and removing. No chemical control was used as there was no pest damage present in the screen house.

#### Set two

Twenty planting pots were filled with sterile soil mixed with manure and sand at the ratio of 3:2:1 and planted with the cowpea seed variety Kunde Tamu as this is the variety from which the pathogen was initially isolated from. DAP fertilizer was applied by placing a pinch of granules on the pot’s soil a week after germination. The plants were watered daily while monitoring the weeds growth and removing. No chemical control was used as there was no pest damage present in the screen house.

#### Pathogen Isolation

Scab symptoms spotted on cowpea variety Kunde tamu planted at farmer demo plots in KALRO Kakamega. Infected pods, stem and leaf were taken to the laboratory for investigation. The pod was rinsed off and air dried. Infected tissue from the pod was then cut off and placed in a 1.5 ml micro-centrifuge tube and 0.2 g/ml of Streptomyces sulphate in 200 ul of distilled water was added. The tissue was macerated with a micro pestle inside the microcentrifuge tube. Full strength PDA media was prepared by mixing commercial PDA 39 g with distilled water and autoclaving for 15 minutes at 121°C at 15 PSI. The media was then let to cool to 40°C and then doped with Streptomyces sulphate at the rate of 1.5g/l as described in the method by (Mungo et al., 1998). The Full strength PDA media doped with streptomycin sulphate (1.5g/l) was poured into a 150 mm sterile petri dish. The macerated infected tissues were then pipettes onto the condensed media at 20 ul pipette drop. A bent glass pipette which was bent by heating the tip narrow end and gently folding it using the wide end of another glass pipette was used to spread the drop across the plate (de Souza et al., 2021). Repeat in three other plates. The plates were monitored regularly and after 7 days a small 1 mm diameter yeast like growth was observed on the plate. The cultures were sub cultured on half strength PDA media plates and let to grow for another five weeks while closely monitoring.

#### Pathogen Identity and Molecular Profile

The identity of the *Elsinoë* sp. isolate used in this study was previously confirmed via multi-locus DNA barcoding of the *ITS* and *TEF1-α* regions and subsequent phylogenetic analysis. The complete molecular characterization, evolutionary lineage, and associated GenBank accession numbers are reported in Oriama et al. (2025)

#### Inoculum preparation

Six weeks old culture plugs were obtained from the petri-dishes by cutting off the plugs from the media. The plugs were then put into 50 ml tubes with distilled water and vortexed vigorously. A haemocytometer was then used to count the conidia present in the extract by putting a drop of the extract on the side of the haemocytometer and letting the extract into the haemocytometer by capillary action. The haemocytometer was then mounted on a light microscope and the scope adjusted to X400 magnification. A motic camera was used to record the observation while counting the conidia.

#### Inoculation

Three weeks old plants both the common beans and the cowpeas were inoculated with the isolated scab. Inoculation was set such that for both sets of the experiments we had a control group and a treatment group. The treatment group was covered in aluminium foil covering the above ground part of the plant in a funnel-like design with an open funnel top that could be closed. Prepared inoculum was then sprayed on the plants from the open funnel top using a plastic pipette at the rate of three sprays and repeated in the evening and the morning after while closing the aluminium foil after inoculation. Watering was done regularly while the plants were let to stay under the aluminium foil for another two days and then transferred to the screen house and set apart from the control group. Monitoring was done to identify any symptoms development on both control and the treatment groups.

#### Monitoring and data collection

Data was collected based on the severity of the scab and the presence of scab symptoms on the plants. Scoring was based on a scale of 1-5, where 1 = 0 % infection (no visible symptoms on either foliage or pods); 2 = less than 10 % scab infection (scattered lesions on either foliage or pods); 3 = 10 to 20 % infection (extensive spotting of young stems and branches or on the pods); 4 = 20 to 50 % infection (stem lesions coalescing, covering half the plants or the pods); and 5 = 50 % or greater infection (foliage severely damaged or pods transformed into mummies containing virtually no seeds), The scores were tabulated and data stored in spreadsheet for downstream analysis.

## RESULTS

Cowpea under field condition displayed scab like symptoms which coalesced to the pod and caused the pod to be mummified with no grain formation and the leaves where folded in a mosaic like manner where the lamina was folded inward starting from the midrib. The stem formed cankers of scab cork-like lesions.

**Figure 1:**
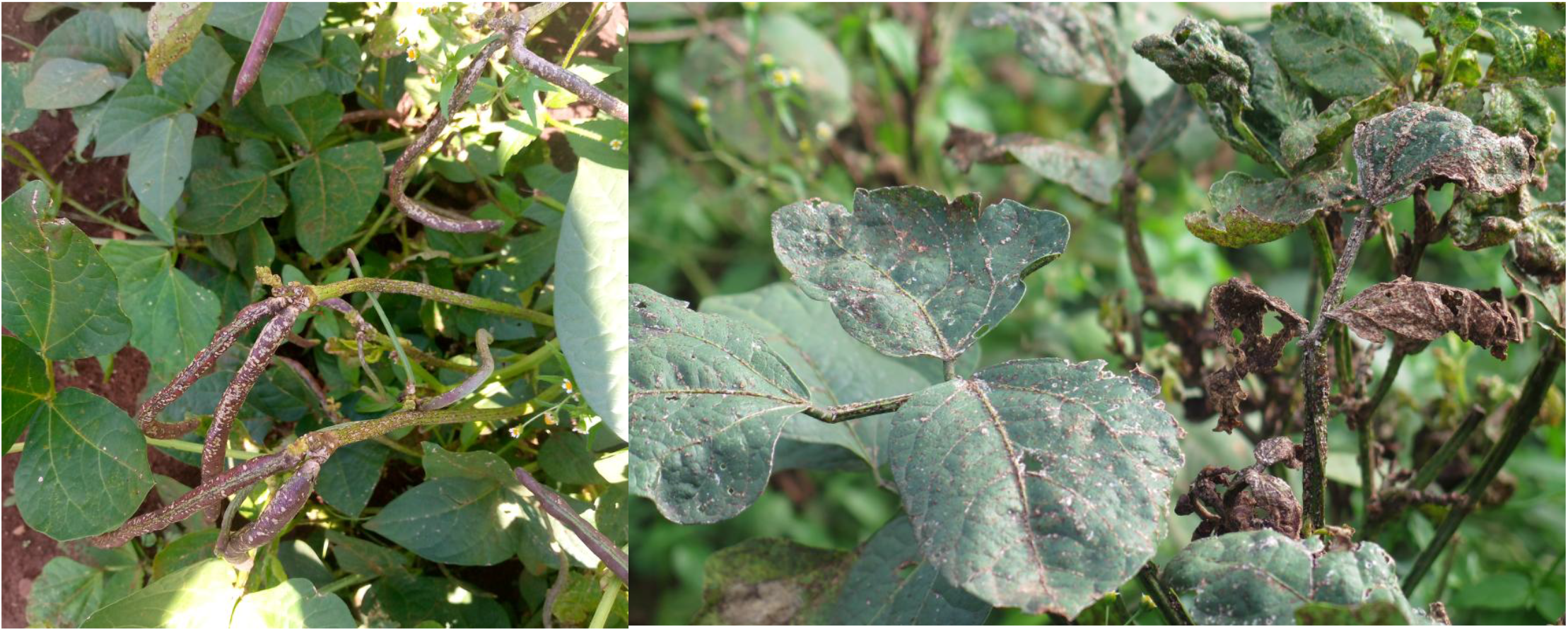
Scab symptoms on cowpea under open field natural infection. Scab lesions on the pod and stem of cowpea plant. Folding of the leave and coalescing of scab lesions on entire plant. Mottle like folding of the leaf.

The colour of the growing scab isolate was cream-yellow and irregularly shaped and branching morphology spreading across the surface with vein-like rough structures which later became surface topology of the radial valley type with undulated edges. The side view was of the crateriform type. The colour gradually changed as the fungi grew to brick red. The texture was rough and the topology was rugged. The fungi had a slow yeast like growing characteristics whereby it had grown 1 mm in 7 days and 3 mm in 14 days. During early growth stages of the fungi

**Figure 2:**
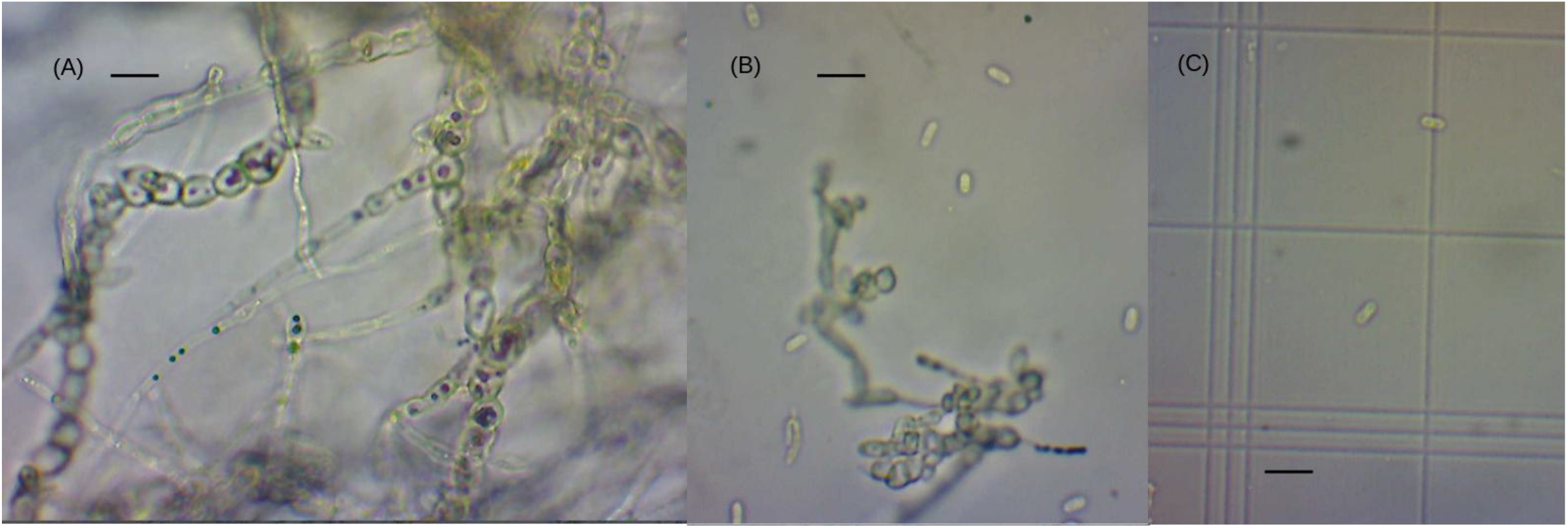
*Elsinoë* spp affecting cowpea viewed under a light microscope at X400 magnification. (A) A dense network of pseudohyphae with small rounded budding-like structure along the pseudohyphae (asexual conidia). (B) pseudohyphae and oval conidia separated and suspended in surrounding medium. Oblong conidia under a counting chamber of haemocytometer with estimates size of 5 um. Bar is 10 um. Source (Oriama et al., 2025) Source: (Oriama et al., 2025)

Under the light microscope the hyphae were pseudo like elongated chain like and were budding. The conidia observed were oblong and non-septate. Oval shaped conidia are seen along the pseudohyphae and are transparent.

**Figure 3:**
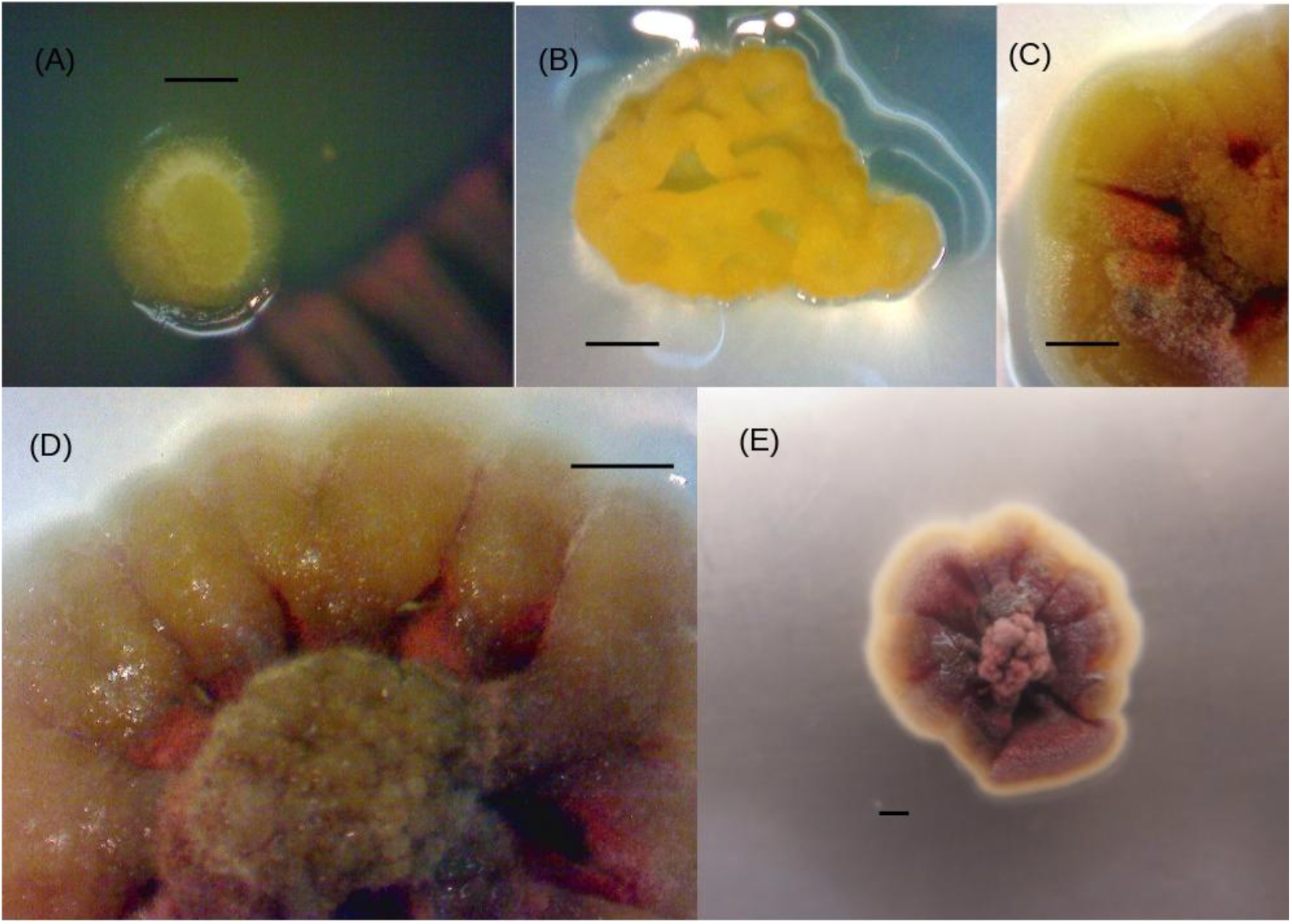
Isolated *Elsinoë* sp morphologies. (A) fungal growth to 1 mm seven days post-isolation on full strength PDA media. (B) Pathogen grows to 3 mm with a distinctive cream yellow colour, clear rugged and vein-like rough topology. (C) At three weeks old the cream yellow colour begins to disappear and a brick red colour begins to sets in. (D) Clear undulated edges of the fungi on PDA media fungi grows to 12 mm at 4 weeks. Source: (Oriama et al., 2025)

Small greying-to-white lesions were spotted on infected plants as early as six days post-inoculation. The lesions spread to cover over 90% of the cowpea leaves by the flowering and pod-filling stages. In contrast, no scab symptoms were observed on common bean plants throughout the growing period. One cowpea variety (Kat Kunde) showed signs of tolerance to cowpea scab, with lesions forming at an early stage but covering less than 10% of the leaves. These symptoms did not progress through its vegetative stage and the flowering stages. The other cowpea varieties, Kunde Tamu and Kunde Soko, were severely affected by the scab. Symptoms appeared on the leaves and stems and later spread to the pods expressing progressive severity up to above 90%. For the pods that formed, most were mummified and failed to fill any grain. Interestingly, at the later pod-filling stages, scab symptoms became severe on the previously observed tolerant variety Kat Kunde, with its pods becoming covered in scab lesions and some mummified due to coalescence of the cork like white lesions and wart like lesions. The treated plants were then transferred and put in the screen house to at meters apart from the control group and left to grow. At two weeks after inoculation the crops developed symptoms such as white spotted lesions on the leaves and three weeks after inoculation the lesions could be seen on the stem. The leaf of the cowpea plants started puckering by folding inward from the lamina like they were mottled. The common bean plants did not show any signs of infection. At week seven post infection the pod of the treatment group formed white raised lesions that where synonymous with scab lesion while the control group did not show any disease symptom. The control group flowered earlier than the treatment group with up to two weeks delay in flowering as the flower bads seemed to be infected with scab for those that emerged. It is worth noting that among the group with several varieties as treatment one variety of cowpea stood out and demonstrated to be tolerant to the disease. The degree of infection was low compared to the other cowpea varieties used in the experiment. This is a commercially released variety.

**Figure 4:**
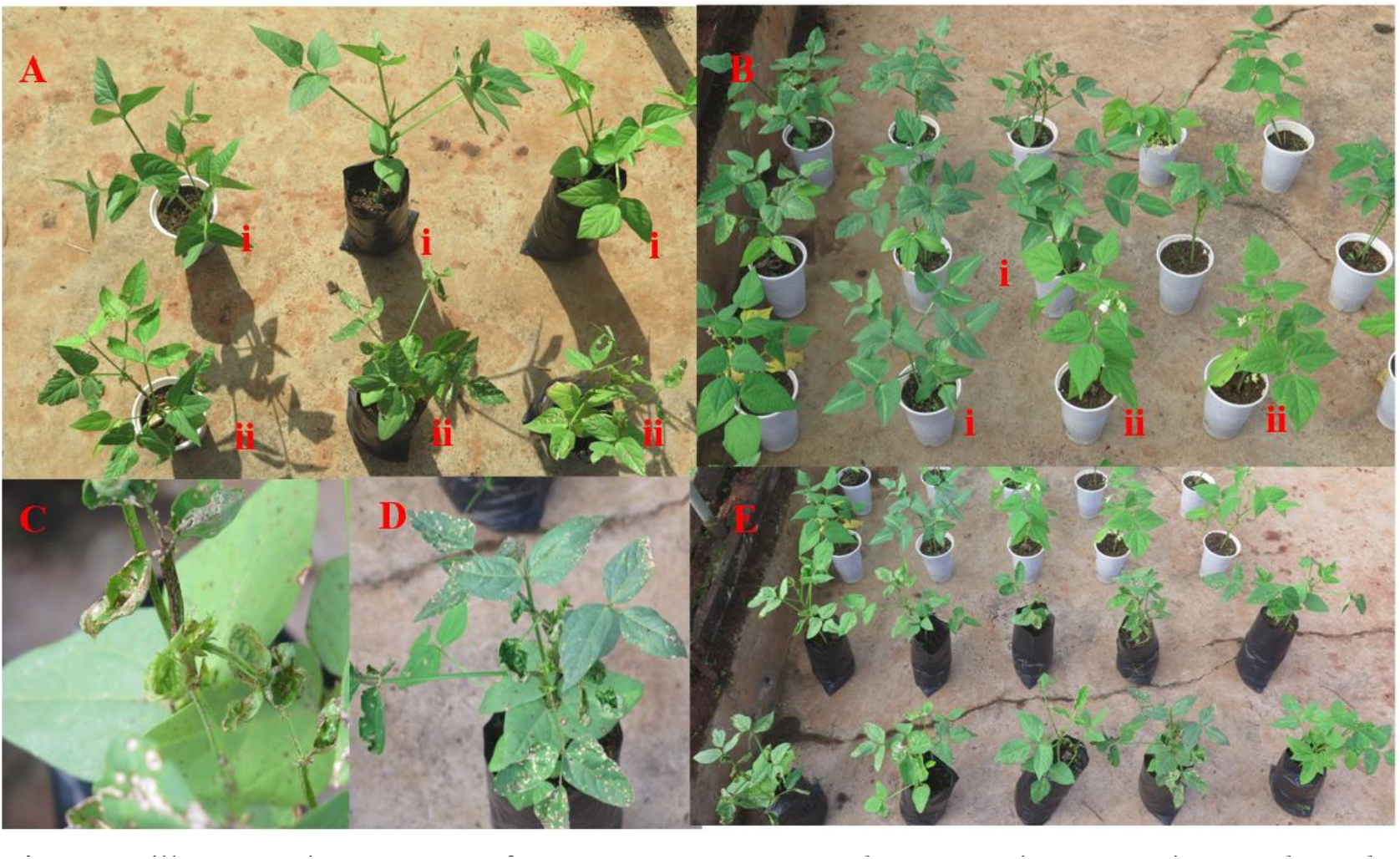
illustrates the response of cowpea (*Vigna unguiculata* L.) and common bean (*Phaseolus vulgaris* L.) varieties at the vegetative stage following inoculation with cowpea scab (*Elsinoë* sp.). **(Ai)** shows the control group of cowpea, while **(Aii)** depicts a sample of the treatment group that was inoculated with cowpea scab, exhibiting early symptoms of infection, including whitish-grey lesions on the leaves and folding of the leaf lamina along the midrib. **(Bi)** highlights a tolerant cowpea variety with minimal symptoms on the leaves, whereas **(Bii)** shows common beans inoculated with cowpea scab that display no signs or symptoms of infection. **(C)** illustrates lesions beginning to coalesce on the stem, and **(D)** shows scab lesions covering more than 10% of the cowpea leaves, accompanied by leaf folding at the midrib. Finally, **(E)** presents the treatment group’s reaction to cowpea scab inoculated on both cowpea and common beans.

One-way analysis of variance showed highly significant differences in scab severity among the legume genotypes (F(5, 18) = 6.851, MSE = 0.931, P = 0.001). Mean separation using the Bonferroni-adjusted LSD test (α = 0.05, critical t = 3.380, df = 18; minimum significant difference = 2.306) placed the genotypes into two overlapping groups (Table 2). The three common bean varieties (CAL 143, CAL 194 and GLP 2) recorded the minimum score of 1 at every growth stage (mean = 1.00), indicating no or negligible symptoms, and were placed in group ‘b’. The cowpea varieties Kunde Soko and Kunde Tamu were the most severely affected, with identical mean scores of 3.50 (group ‘a’). Kat Kunde had a lower mean of 2.75, scoring 2 from 6 DPI through flowering before rising to 5 at pod filling. Because its difference from the other cowpeas (0.75) was smaller than the MSD (2.306), and it also did not differ significantly from the common beans, it was placed in the intermediate group ‘ab’ and is best described as moderately tolerant rather than resistant.

**Table 1:** Plant materials used in the Koch’s postulate and cross infectivity experiment.

| Cowpea (*Vigna unguiculata*. L) | Common Bean (*Phaseolus vulgaris*. L) |
| --- | --- |
| Kunde tamu. | GLP 2 |
| Kunde soko | CAL 143 |
| Kat kunde | CAL 194 |

**Table 2:** Severity scores post inoculation with isolated pathogen across different growing stages of both cowpea and common bean.

| Crop | Crop Variety | Six DPI | Vegetative stage | Flowering stage | Pod Filling Stage | Mean Severity |
| --- | --- | --- | --- | --- | --- | --- |
| Cowpea | Kunde Tamu | 2 | 3 | 4 | 5 | 3.50 a |
| Cowpea | Kunde Soko | 2 | 3 | 4 | 5 | 3.50 a |
| Cowpea | Kat Kunde | 2 | 2 | 2 | 5 | 2.75 ab |
| Common Bean | CAL 194 | 1 | 1 | 1 | 1 | 1.00 b |
| Common Bean | CAL 143 | 1 | 1 | 1 | 1 | 1.00 b |
| Common Bean | GLP 2 | 1 | 1 | 1 | 1 | 1.00 b |

Means followed by the same letter are not significantly different (Bonferroni-adjusted LSD, $\alpha = 0.05$ ; MSD = 2.306).

**Figure 5:**
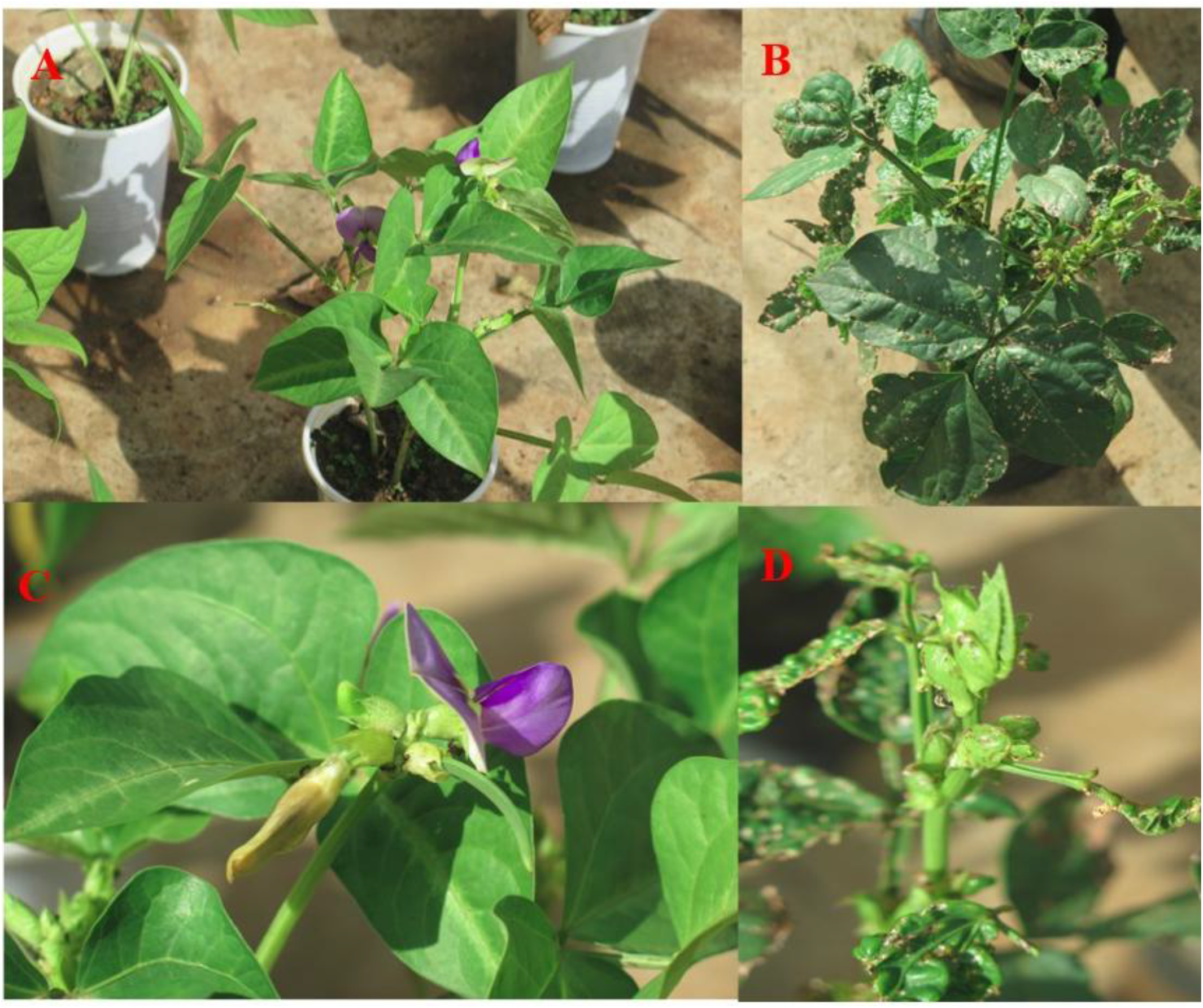
The effects of cowpea scab on cowpea plants at the flowering stage. **(A)** and **(B)** show a sample of the control group with unaffected leaves and flowers, flowering normally. In contrast, **(C)** and **(D)** depict the treatment group, which is severely affected by cowpea scab, resulting in the failure of bud formation and flower abortion.

**Figure 6:**
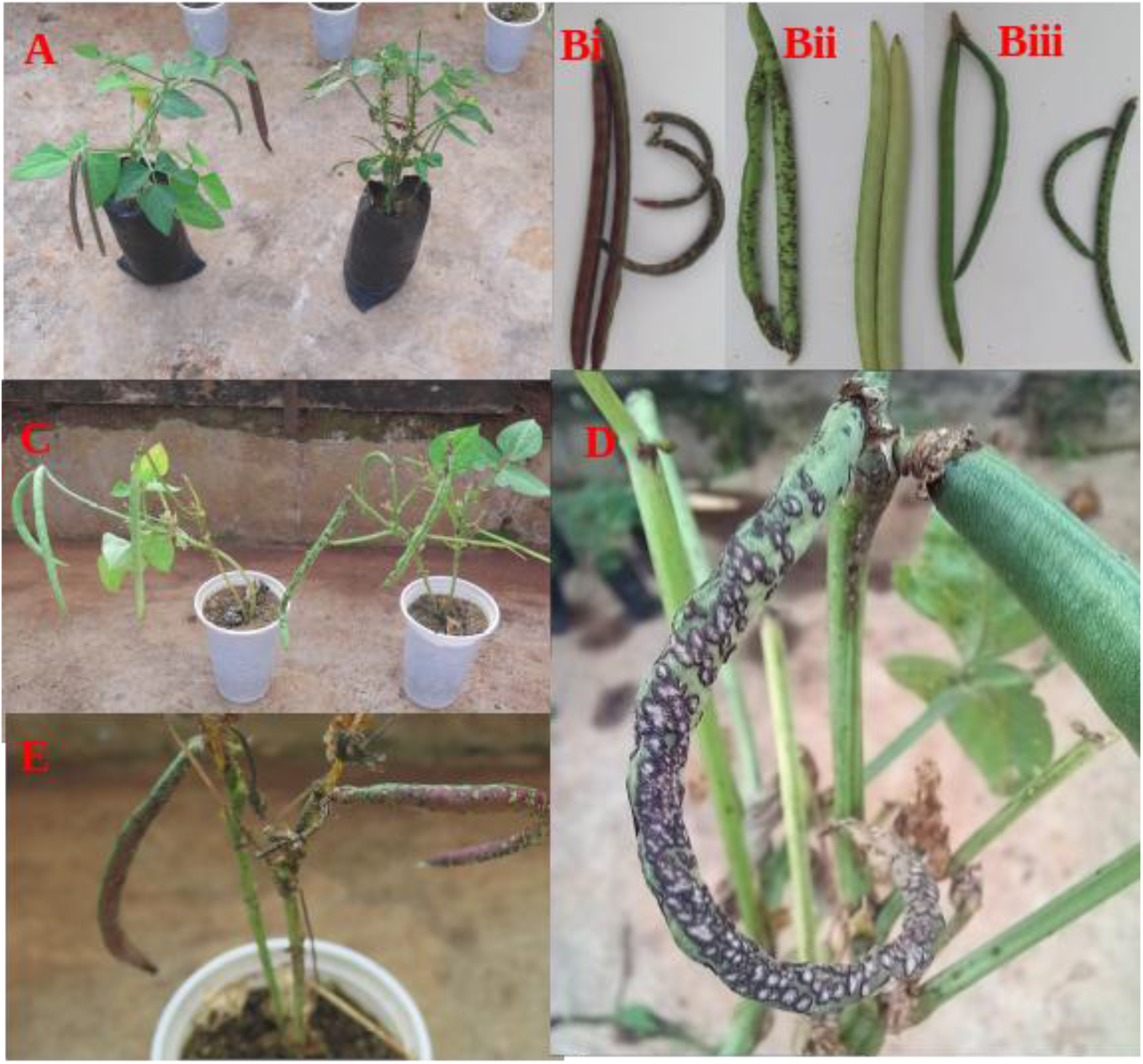
Cowpea plants at the pod-filling stage while infected with cowpea scab. **(A)** compares the control and treatment groups, with the treatment group being severely affected, resulting in the failure of pod formation. **(Bi), (Bii)**, and **(Biii)** show the healthy control group with well-filled pods, while the treatment group exhibits pods covered in scab lesions, folding, and poor pod filling. **(C)** highlights cowpea plants that reached the pod-filling stage despite being infected with cowpea scab. **(D)** and **(E)** display infected pods, with one pod mummified due to the severity of the scab infection.

## DISCUSSION

This work forms part of the first report of *Elsinoë* spp. causing Cowpea scab in Kenya (Oriama et al., 2025). Cowpea scab caused by *Elsinoë* spp. is shown to be a highly destructive disease under both natural field conditions and controlled inoculation. The observed symptoms including cork-like lesions on leaves, stems, and pods, severe leaf puckering, and pod mummification are consistent with previous descriptions of scab diseases in legumes (Emechebe, 1980; Fan et al., 2017; Masheti et al., 2024; Oriama et al., 2025). The coalescing nature of lesions leading to extensive tissue damage and distortion highlights the aggressive pathogenicity of the isolate used in this study.

Under field conditions, the disease caused severe deformation of vegetative and reproductive tissues, particularly the inward folding of the leaf lamina along the midrib and the formation of cankers on stems. These symptoms significantly reduce the photosynthetic capacity of the plant and disrupt normal physiological processes (Oliver, 2024). The mummification of pods and absence of grain formation observed in heavily infected plants further demonstrate the direct impact of the pathogen on yield. Similar findings have been reported in other Elsinoë-induced scab diseases, where infection during reproductive stages leads to substantial economic losses due to reduced marketability and yield (Masheti et al., 2024b; Pham et al., 2024).

The cultural and morphological characteristics of the isolated pathogen corroborate its identification as *Elsinoë* spp (Fan et al., 2017). The slow-growing, yeast-like colonies, transitioning from cream-yellow to brick red with rugged and crateriform morphology, align with descriptions of Elsinoë species in previous studies. Microscopic observations of pseudohyphae with budding, non-septate, oblong conidia further support this classification. The presence of pseudohyphal structures suggests an adaptation that may enhance survival and infection efficiency under varying environmental conditions.

A key finding of this study is the apparent host specificity of the pathogen. Despite successful infection and severe symptom development in cowpea, no symptoms were observed in common bean (*Phaseolus vulgaris*) across all growth stages. This strongly supports the hypothesis that *Elsinoë* spp. exhibit host specialization, with limited or no cross-infectivity between cowpea and common beans. Such specificity has important implications for disease epidemiology and management, as it suggests that cowpea scab pathogens may not pose a direct threat to common bean production, even in mixed cropping systems.

The progression of disease severity observed across growth stages indicates that cowpea is most vulnerable during the flowering and pod-filling stages. Early infection, as observed at six days post-inoculation, rapidly escalated to severe disease covering more than 90% of leaf area in susceptible varieties. The delay in flowering and the occurrence of flower abortion in infected plants further emphasize the impact of the pathogen on reproductive development. These findings suggest that infection during early vegetative stages predisposes the plant to more severe outcomes later in its growth cycle.

Variation in varietal response to scab infection was also evident. While Kunde Tamu and Kunde Soko were highly susceptible, exhibiting progressive disease severity culminating in severe pod damage and yield loss, Kat Kunde initially demonstrated tolerance, with limited lesion development during vegetative and flowering stages. However, this tolerance was not sustained, as severe symptoms developed during the pod-filling stage. This indicates that resistance in Kat Kunde may be partial or stage-dependent rather than complete. The common bean varieties (CAL 194, CAL 143, and GLP 2) remained completely unaffected, reinforcing the concept of host specificity.

The identification of a tolerant cowpea variety, even with late-stage susceptibility, is of significant importance for breeding programs. Partial resistance can still be valuable in integrated disease management, particularly if combined with other control strategies. However, the breakdown of tolerance at later stages underscores the need for screening across the entire growth cycle when evaluating resistance to cowpea scab.

The findings of this study highlight the severity of cowpea scab and its potential to cause substantial yield and economic losses. The clear host specificity observed suggests that management strategies should focus primarily on cowpea, without immediate concern for cross-infection to common bean. Furthermore, the variability in varietal response provides a foundation for future breeding efforts aimed at developing durable resistance. Additional studies incorporating molecular characterization of the pathogen and resistance mechanisms in cowpea would further enhance understanding and support the development of effective control strategies.

## CONCLUSION

Cowpea scab infection is specific to cowpea. There are disease resistant cowpea varieties in existence since with a small sample population we observed tolerance on Kat kunde variety.

## RECOMMENDATIONS

Further research should focus on breeding for durable resistance and understanding the genetic and pathogenic variability of *Elsinoë* spp. to support sustainable disease management strategies

## ACKNOWLEDGEMENT

A version of this article was presented at the 2nd KALRO Scientific Conference and Innovation Expo 2026 (15th–19th June 2026, Nairobi, Kenya) and has been submitted to the conference proceedings for publication.

The authors gratefully acknowledge the Kenya Agricultural and Livestock Research Organization (KALRO), Kakamega, for facilitating this research through provision of field space, materials, and technical support.

## Notes

### Competing Interest Statement

The authors have declared no competing interest.

